# Development shapes molecular responses to thermal extremes in a desert bird

**DOI:** 10.64898/2026.05.29.728694

**Authors:** Lea L. Huber, Charlie K. Cornwallis, Molatelo R. Kekana, Neleke Lotz, Zanell Brand, Schalk Cloete, Anel Engelbrecht, Mads F. Schou

## Abstract

Thermal extremes are among the most immediate environmental challenges faced by animals. Coping with these conditions across life is complex because growth changes body size, heat exchange and thermoregulatory demands. Consequently, adaptive responses in young, small individuals may be maladaptive, or require adjustment, later in life. However, in endotherms we know very little about how thermoregulatory responses to hot and cold temperatures change during development, or about the molecular mechanisms regulating responses. We examined the molecular responses to acute heat (40°C), cold (12°C) and benign (23°C) conditions in 1- and 8-week-old ostrich chicks (*Struthio camelus*), a rapidly growing species exposed to strong daily and seasonal temperature variation. We found that in 1-week-old chicks, 44% of temperature-related genes were involved in both heat and cold responses, indicating that responses to opposing temperatures involve overlapping molecular pathways. However, the response of 76% of these genes changed during development. For example, some genes that increased with heat exposure when young decreased with heat exposure later in development, and vice versa. Such opposing selection pressures may maintain genetic variation in thermoregulatory pathways. Consistent with this prediction, comparisons between ostrich subspecies adapted to different thermal environments revealed patterns of genomic variation compatible with balancing selection in differentially expressed genes. Our results show that hot and cold temperatures trigger overlapping molecular responses that change during development, which shape genetic variation in the thermoregulatory system.

## Introduction

Maintaining internal body temperatures while external conditions fluctuate is critical for survival in endotherms. In a warming climate, individuals must adapt to more frequent and intense heat waves while also coping with cold temperatures (Perkins-Kirkpatrick and Lewis, 2020; Fischer, Sippel and Knutti, 2021), especially in regions with great seasonal and daily temperature fluctuations (Bathiany *et al*., 2018). This may be challenging because heat and cold tolerance are not necessarily independent, and can be negatively correlated, thereby limiting adaptation (Rodríguez-Verdugo *et al*., 2014; Sørensen *et al*., 2016; Schou *et al*., 2022). The impact of fluctuating temperatures on individuals is also predicted to change during development, where thermoregulatory demands and capacities shift with growth. Consequently, animals must continually balance their investment in heat and cold tolerance throughout development, but the extent to which endotherms do this, and the molecular mechanisms involved, are poorly characterised.

During growth, an individual’s surface-area-to-volume ratio typically decreases, reducing the capacity to dissipate metabolic heat via the surface (Rowe *et al*., 2013; Riddell *et al*., 2019). Consequently, larger animals face greater difficulty thermoregulating under sustained heat, whereas smaller animals struggle to retain heat in the cold (Scholander *et al*., 1950). Thermoregulatory demands are therefore predicted to shift throughout ontogeny, especially in large endotherms that undergo substantial changes in body size during development (Rotenberry and Balasubramaniam, 2020). However, most research has focused on a single developmental stage, and studies have rarely integrated responses to both heat and cold exposure. As a result, remarkably little is known about how thermoregulatory mechanisms operate across life stages (Schou and Cornwallis, 2024). This represents a significant gap, as population persistence critically depends on thermal tolerance throughout life (Dahlke *et al*., 2020).

Understanding how thermoregulatory mechanisms change across life stages requires identifying the underlying molecular processes that enable physiological adjustment. In endotherms, changes in gene expression are central to regulating physiological responses to environmental conditions, but few studies have tested whether the same genes are involved in heat and cold responses at different developmental stages (Monson *et al*., 2018; Kim *et al*., 2021; Ballinger *et al*., 2023). Several core mechanisms, such as metabolic rate and vasodilation, contribute to both warming and cooling responses, and at least part of the molecular response to heat and cold is expected to be shared.

Shifting thermoregulatory demands during ontogeny could alter selection on genes regulating heat and cold responses across life stages. This may favour the evolution of gene expression patterns that optimise thermoregulatory performance during each life stage. Alternatively, age-related changes in gene expression may only partly resolve thermoregulatory conflicts between life stages, resulting in opposing directional selection on the same genes at different ages. For example, in Drosophila, genes involved in starvation resistance are under opposing directional selection pressures at different ages (Kawecki *et al*., 2021). Opposing selection pressures across life stages can generate balancing selection, maintaining diversity in genes, but may also constrain adaptation to fluctuating temperatures.

Here we investigate the molecular basis of thermoregulation in the common ostrich (*Struthio camelus*) during development. Ostriches are native to highly fluctuating thermal environments across Africa (-5°C to 50°C) and undergo one of the largest changes in body size during development of any endotherm (∼0.7 to 100 kg), potentially making ontogenetic shifts in thermal responses pronounced in this species. Ostrich chicks are highly precocial, leaving their nest and foraging within hours of hatching, which means they are immediately exposed to ambient temperatures. Parents can, however, protect their chicks from both cold nights and hot days by brooding them up and providing shade until around six weeks of age. This is especially important in their southern range, where temperature fluctuations are greater and nights are colder than in eastern Africa (Fick and Hijmans, 2017; Svensson *et al*., 2023). We exposed 1- and 8-week-old ostriches to three manipulated temperature environments (mean temperatures: cold = 12°C, benign = 23°C, hot = 40°C, Fig 1A-C) and quantified their gene expression responses in blood samples. We then tested: 1) whether the molecular responses to hot and cold temperatures have a shared genetic basis; 2) how the molecular thermal response is modulated across age; and 3) whether thermoregulatory genes whose response changes with age show signatures of balancing selection in two ostrich subspecies native to different climates.

**Figure 1:**
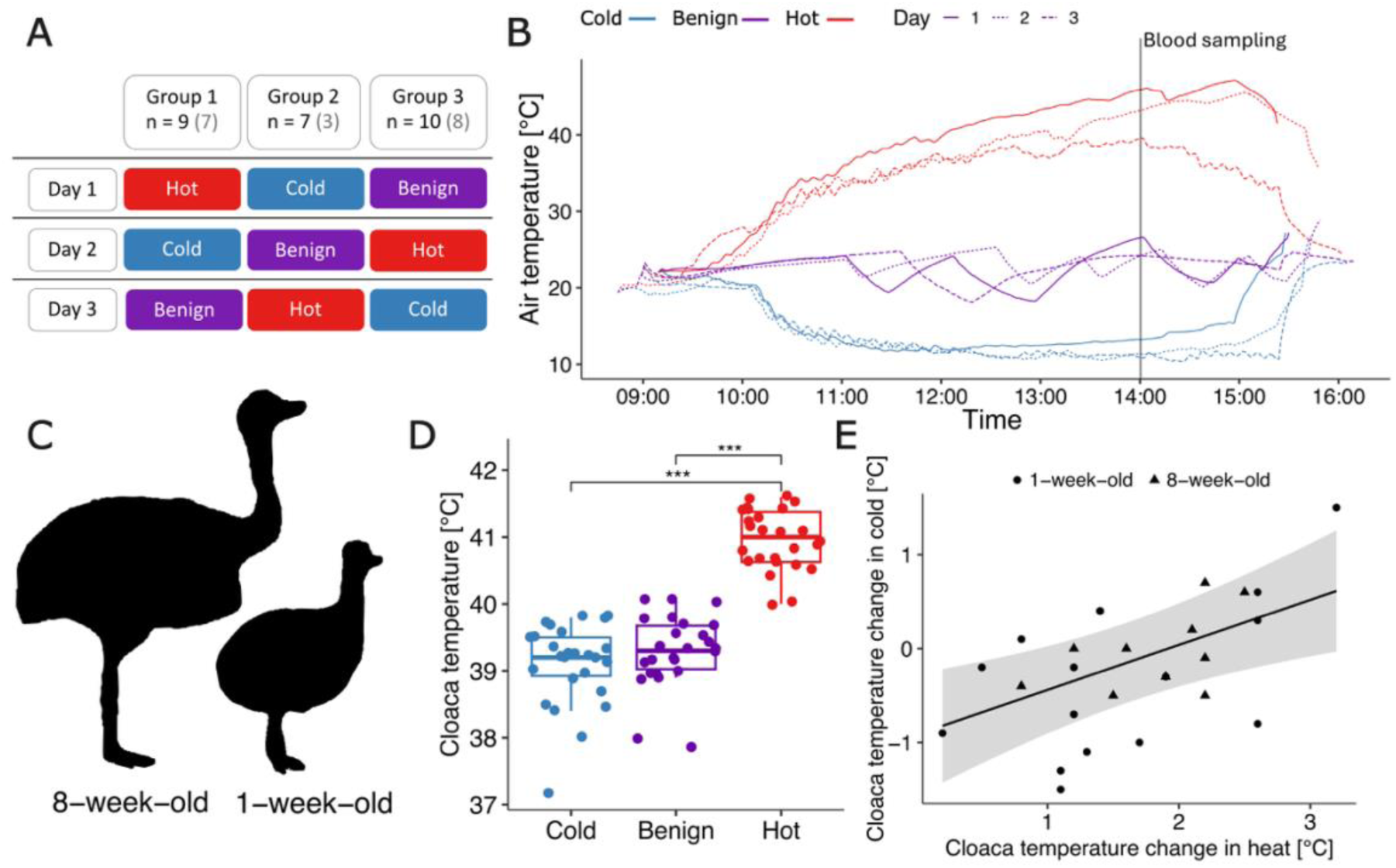
Experimental manipulation of acute temperature exposure of ostrich chicks during development. A) Illustration of the experimental design with three groups exposed to benign, hot and cold air temperatures in a randomised order across three days. Numbers in black refer to the number of chicks whose cloacal temperature was measured on each day, numbers in grey refer to the number of chicks whose transcriptome was analysed. B) Temperature exposure took place in experimental chambers where chicks were placed at ∼09:00 in the morning. The temperature was then gradually increased (hot), maintained (benign), or decreased (cold) until blood sampling commenced at 14:00 (vertical line). C) N = 16 of the chicks acutely exposed to heat, cold and benign temperatures were 1-week-old hatchlings (9 for transcriptome analyses), and n = 10 were 8-week-old chicks (9 for transcriptome analyses), whose typical size difference is illustrated with silhouettes. D) After blood sampling, the cloacal temperature was measured as an indication of body temperature change during the temperature treatment (see Table S1 for coefficients and p-values). E) Individuals that avoided reductions in body temperature under cold conditions had higher body temperatures when exposed to hot conditions (approximated by cloacal temperature). Each point represents the individual change in body temperature in the cold (cold – benign) versus in the heat (heat – benign). The fitted line was estimated with a linear model, and the shaded area represents the 95% confidence interval (see Table S2 for coefficients and p-values).

## Results

### The regulatory response to heat is greater than to cold

We analysed gene expression count data using generalised linear mixed models, with the three temperature treatments, age and sex as categorical variables and correcting for individual and group effects. The expression of 3.8% of all genes was influenced by temperature (post hoc tests: 416 out of 11075 genes at q-value 0.01). About half of these genes (214 genes, 51.4%) responded to either heat or cold at 1- and 8-weeks of age (see also “distinct response” in Fig 2A). Of these genes, most were differentially expressed in the heat treatment compared to the benign treatment (91.6%), with only 8.4% differentially expressed in the cold compared to benign conditions. This was in accordance with the cloacal temperatures being similar in the cold and benign treatments (Fig 1D). Cloacal temperature may have been buffered by behavioural responses, with huddling and shivering being observed in the cold treatments.

**Figure 2:**
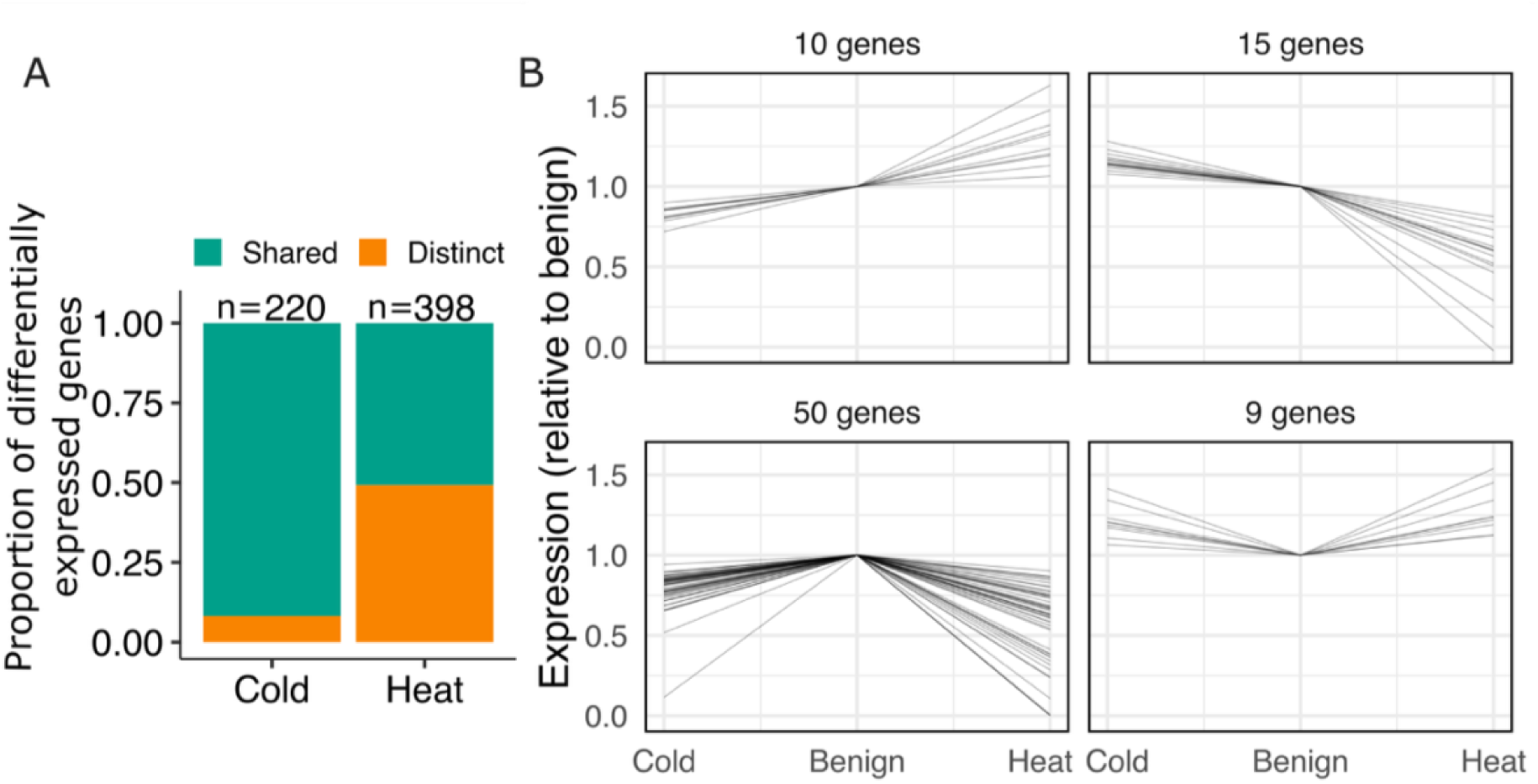
Expression patterns of genes responding to heat and cold. A) Proportion of genes that were differentially expressed during heat or cold exposure (distinct response) versus genes that responded to both heat and cold exposure (shared response). B) Standardised expression patterns of genes responding to both heat and cold exposure (shared response) in the same way across both age groups. Genes were classified into having distinct roles, being downregulated in cold and upregulated in heat (10 genes) or upregulated in cold and downregulated in heat (15 genes), or showing a general stress response, being downregulated in cold and heat (52 genes) or upregulated in cold and heat (8 genes). Differential expression in heat was generally greater than that in cold compared to the benign treatment.

### Genetic responses to heat and cold are partly shared

A substantial proportion of differentially expressed genes responded to both heat and cold in 1-week-olds (193 genes, 46%) and in 8-week-olds (147 genes, 35%), resulting in 202 genes differentially expressed in both heat and cold in either or both age groups (48.6%). It is notable that of all genes differentially expressed in response to cold, 92% were also affected by heat. Conversely, of the genes that responded to heat exposure, only 51% were affected by cold (Fig 2A).

Genes involved in thermoregulation may respond similarly to both heat and cold, indicating a general stress response, or in opposite directions, indicating distinct roles in warming and cooling. To distinguish between these responses, we focused on the 84 genes that were consistently involved in responses to both heat and cold across both age groups (20.2% of genes, Fig 2B). Most of these genes were consistently upregulated in response to both heat and cold exposure or consistently downregulated in response to both heat and cold exposure (59 genes; Fig 2B, bottom panels), in accordance with a general acute temperature stress response. However, a substantial subset of genes (25 genes, 30%) showed opposing expression patterns, being upregulated in heat and downregulated in cold or vice versa (Fig 2B), suggesting distinct roles in thermoregulation during heat and cold exposure. It therefore appears that heat and cold responses have a partly shared genetic basis, which could constrain their capacity to evolve independently. In line with this, individuals more resistant to cold were less resistant to heat (linear model of change in cloacal temperature: F_(2,18)_ = 11.1, p-value = 0.004, Fig 1E, Table S2).

### Expression of thermoregulatory genes changes during development

We found 377 genes whose response to acute temperature stress was modulated by age, either as a change in the direction or intensity of the temperature response across age, or as a temperature response only present in one age group (post hoc tests of age by temperature interaction; q-value < 0.01). There was a large overlap between genes involved in the shared response to heat and cold, and genes that changed expression with age. For example, 76% of shared response genes in 1-week-olds had a changed expression in 8-week-olds.

For some of the age-modulated genes (heat: 25.5%/96 genes; cold: 9.8%/37 genes), the pattern of expression was the same across ages, but the intensity of expression changed (Fig 3B-C “same direction”). For most of these genes, the intensity of the response was higher in 1-week-olds than in 8-week-olds (Fig 3B-C). Other genes showed a change in the direction of regulation, for example, the response to heat changed from upregulation to downregulation between ages. We found such a pattern of expression in 12% of age-responsive genes (46 genes, Fig 3B-C “opposite direction”), with most genes being downregulated in 1-week-olds and upregulated in 8-week-olds in cold (scatterplot in Fig 3B, upper left quadrant), or upregulated in 1-week-olds and downregulated in 8-week-olds in heat (scatterplot in Fig 3C, lower right quadrant). We also identified several genes that were differentially expressed only in one age group (Fig 3B-C, see also Table S3).

**Figure 3:**
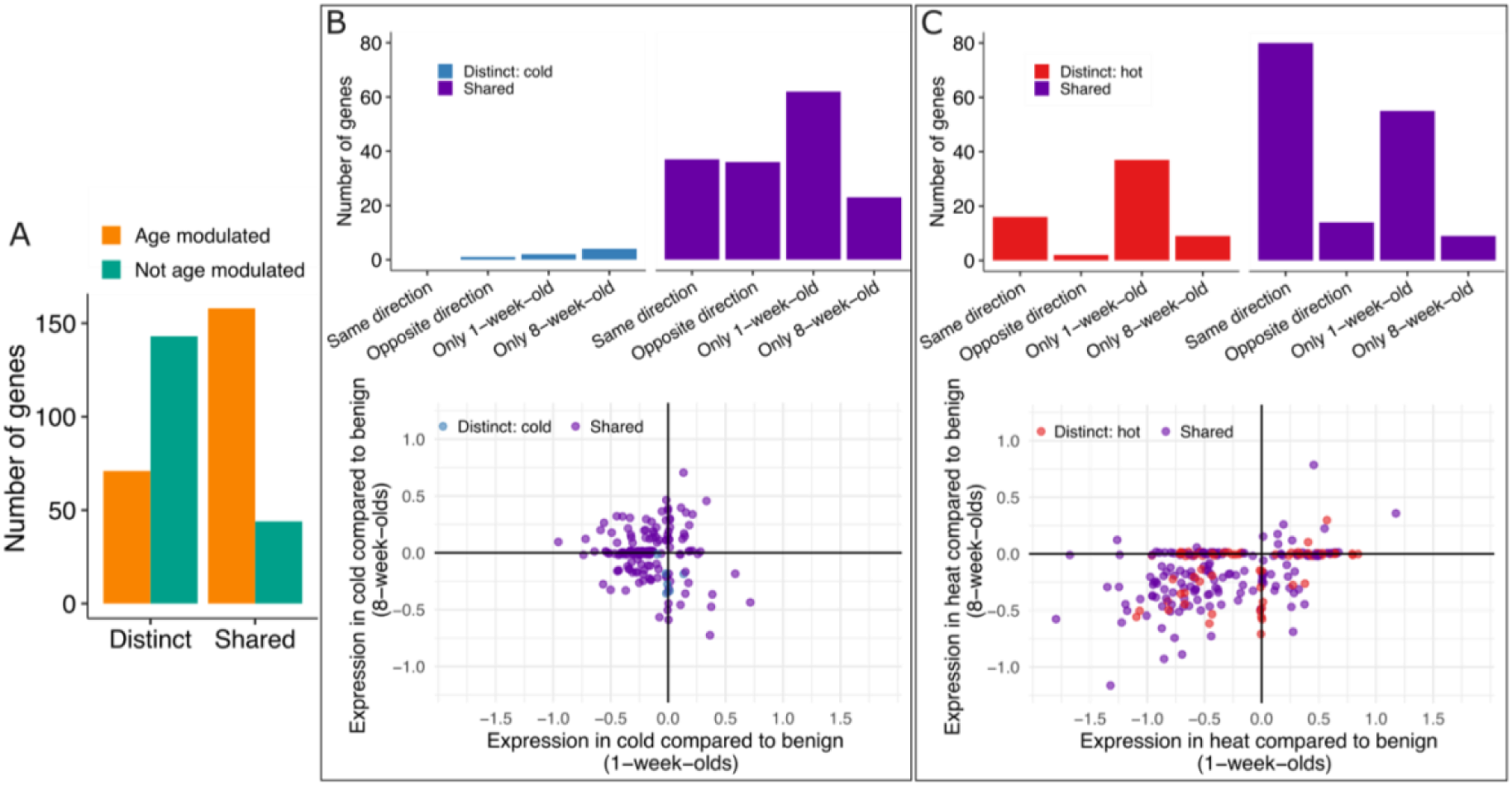
Genes modulated by age were more frequently involved in both heat and cold responses. A) The majority of genes responding to both heat and cold exposure (shared response) did also change expression across ages. In contrast, most genes only involved in responses to either heat or cold (distinct response) were unaffected by age. B-C) The genes affected by age changed their expression in different ways: i) same direction – genes were for example upregulated in both age groups but by different amounts (upper right and lower left quadrants); ii) opposite direction – genes upregulated in one age group were downregulated in the other or vice versa (upper left and lower right quadrants); iii) only 1-week-old – genes were only differentially expressed in 1-week-olds (on horizontal line); iv) only 8-week-old – genes were only differentially expressed in 8-week olds (on vertical line). The bar plots show the number of genes in each of these categories for distinct response and shared response genes separately, in cold (B) and heat (C). The scatterplots show estimated effects of heat and cold on gene-expression in age-modulated genes extracted from a generalized linear mixed model. Genes that were only differentially expressed in heat across ages are shown in red, genes that were differentially expressed in both treatments at any age are shown in purple, and genes only differentially expressed in the cold treatment across ages are shown in blue. Points are positioned with a 0.02 jitter. See Fig S3 for a scatterplot of genes not modulated by age.

### The shared temperature response is a general stress response

To determine whether genes responding to either heat or cold exposure are functionally distinct from genes responding to both conditions, we performed GO-term enrichment and KEGG analyses. Genes not modulated by age and only upregulated in heat were enriched for functions mostly related to protecting proteins from denaturation during thermal stress (e.g. GO-terms: response to unfolded protein, protein folding, KEGG: spliceosome, Table S4), whereas genes only upregulated in the cold treatment were mostly involved in adjusting lipid metabolism (e.g. GO-terms: (cellular/glycero-)lipid metabolic process, KEGG: glycerolipid metabolism, Table S4). In contrast, genes that responded to both heat and cold exposure had more diverse functions, which were related to the immune system, inflammatory responses, metabolism, cell signalling and development (Table S4). This suggests a more general response to organismal damage or departure from homeostasis, which may be triggered by a variety of stressors, such as disease and temperature, that were activated during both heat and cold exposure.

The functional role of genes that responded differently to heat or cold exposure across age groups was also diverse, with multiple overrepresented terms only in genes exclusively expressed in 8-weeks-olds and in genes changing direction of expression. Their functions included external stimuli, cell adhesion, and signalling in heat response genes, while in cold response genes, functions included endocytosis, apoptosis and potassium ion transport (Table S5).

### Temperature related genes under selection in ostrich subspecies

Genes that shift the direction of expression in response to acute temperature stress across ages may be under opposing directional selection at different life stages, resulting in balancing selection that maintains nucleotide diversity. We examined this possibility by whole-genome sequencing multiple individuals from two ostrich subspecies, the Masai ostrich (*S. c. massaicus,* n = 6) and the South African ostrich (*S. c. australis*, n = 8) that are distinct from the farmed population used for the experiment, to estimate within subspecies nucleotide diversity. We found elevated nucleotide diversity in the set of genes that had opposing expression patterns across ages compared with genes not modulated by age, consistent with hypothesised balancing selection (Fig 4A).

**Figure 4:**
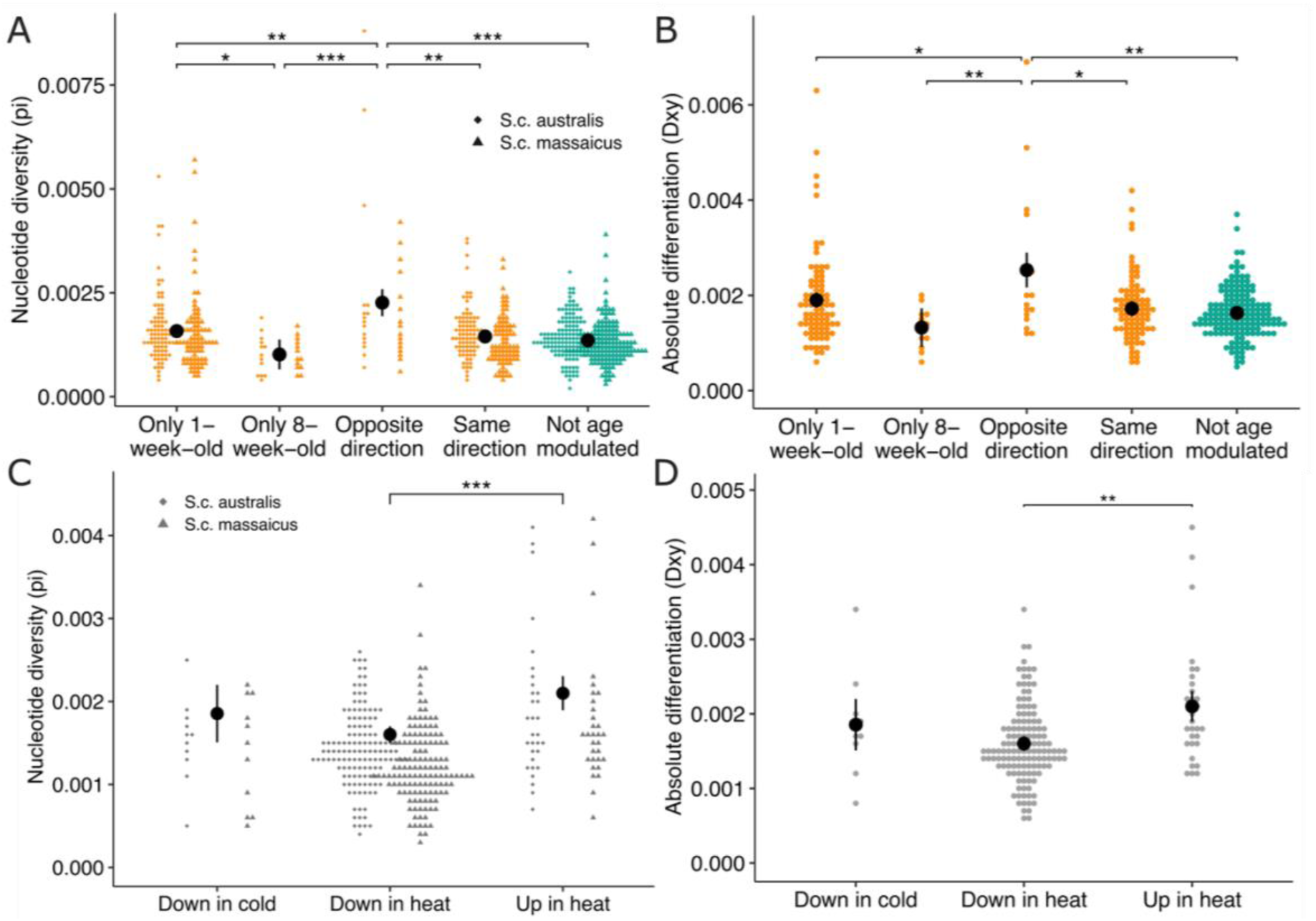
Genes involved in thermoregulation show signals of balancing selection across ostrich subspecies. Nucleotide diversity (pi) and absolute differentiation (D_xy_) of S.c. australis and S.c. massaicus are higher in genes with opposing expression patterns across ages, as well as in genes upregulated in response to heat regardless of age. Only genes with > 10 SNPs and only gene sets with > 10 genes were included in the analyses. (A-B) Pi and D_xy_ of genes responding differently to temperature exposure between 1-week and 8-week-old ostrich chicks. Age-modulated genes (in orange) were grouped into genes expressed in only one age group, in opposite directions across age groups at one temperature (e.g. upregulated in 1-week-olds and downregulated in 8-week-olds in heat) and expressed in the same direction across age groups at one temperature (e.g. upregulated in both 1-week-olds and 8-week-olds in heat but by different intensities). Genes not modulated by age were used for control comparisons (“Not age modulated” in green). (C-D) Pi and D_xy_ of genes only responding to either heat or cold exposure in both age groups (distinct). Genes are grouped into genes downregulated in the cold treatment compared to the benign treatment (Down in cold) and genes downregulated (Down in heat) or upregulated (Up in heat) in the heat treatment. Black dots are least-square means determined with linear mixed models for pi estimates and linear models for Dxy estmates, bars are 95% confidence intervals. Significant differences between categories were determined using pairwise comparisons and adjusted with Tukey’s method. See Fig S7 A-B for pi and D_xy_ in genes with a shared response in heat and cold exposure.

The Masai ostrich originates from eastern Africa, which has an overall warmer and less seasonal climate that has likely resulted in elevated heat tolerance compared to the South African ostrich (Schou *et al*., 2022; Svensson *et al*., 2023). We therefore tested for signatures of genomic differentiation in genes responding to temperature across these two the ostrich subspecies. We found an elevated absolute divergence (D_xy_) between subspecies in genes that changed the direction of expression in response to temperature across age groups (Fig 4B) but limited relative divergence (F_ST_) (Fig S8). F_ST_ is influenced by diversity within populations, and thus it is expected to be reduced in the same genes that show elevated nucleotide diversity. In contrast, D_xy_ is not affected by current levels of nucleotide diversity, but it can be affected by ancestral diversity (Cruickshank and Hahn, 2014). Elevated nucleotide diversity and D_xy_ are consistent with these genes being under balancing selection within subspecies and differentiating across subspecies due to their different climatic origins. We found similar evidence for balancing selection in genes upregulated in heat that were not modulated by age (Fig 4C-D, see Fig S9 for genomic analyses corrected for the effects of gene length). Despite low sample sizes limiting possibilities for further analyses of site frequency spectrum in these genes, our results indicate that several sets of genes involved in thermoregulation may be under balancing selection and diverging across natural populations.

## Discussion

Coping with temperature fluctuations early in life, before physiological systems have fully developed, can be challenging for endotherms. We found that a substantial proportion of genes differentially expressed in response to acute temperature exposure were involved in both heat and cold responses, and that these shared genes were more likely to change expression with age. This indicates that development after hatching requires modulation of the combined thermal response, consistent with the expectation that relative sensitivity to heat and cold shifts throughout development.

The shared molecular response to heat and cold appears to involve genes involved in general homeostasis, as their functions are related to metabolism, cell signalling and hormone biosynthesis, amongst others. Thermal responses also activated immune-related genes, which is consistent with findings in ectotherms (Yin *et al*., 2025), and suggests that departures from physiological homeostasis may increase vulnerability to pathogens (Tang *et al*., 2021; Liu *et al*., 2025). Generally, genes involved in multiple functions that are under different selection pressures can have limited potential to evolve (Chen and Zhang, 2020). Likewise, genes expressed across thermal extremes could have reduced evolutionary potential in fluctuating environments as mutations beneficial for coping with heat may be detrimental for coping with cold and vice versa. This is supported by the finding that chicks whose cloacal temperature was more affected by cold were less affected by heat. Furthermore, previous research in ostriches showed a negative correlation between individual egg laying rates in hot and cold conditions, indicating a trade-off between heat and cold tolerance (Schou *et al*., 2022).

Heat exposure triggered a distinct molecular response from cold exposure, which involved the upregulation of several heat shock proteins (HSPA2, HSPA5, HSPA8, HSPB9, HSPD1, DNAJA4, HSP90AA1, and its interaction partner AHSA1). HSPs act as chaperones in protein stabilisation and (re-)folding (Gething and Sambrook, 1992), and are involved in heat responses in organisms across the tree of life, including birds (Craig and Schlesinger, 1985; Xie, Tearle and McWhorter, 2018). In aquatic ectotherms, HSPs have been reported to be under balancing selection, with different variants thought to confer advantages under specific thermal conditions (Chen *et al*., 2017; Junprung *et al*., 2021). Our findings are consistent with these results: genes upregulated in heat showed both high absolute differentiation across two ostrich subspecies, a sign of local adaptation to the thermal environments of eastern and southern Africa, and high nucleotide diversity, suggesting that additional forces, such as balancing selection, preserve genetic variation within populations.

A subset of age-modulated genes (11%, 46 genes) reversed the direction of regulation between age groups. Functionally, these age- and temperature-dependent genes were related to synaptic vesicle endocytosis and potassium ion transport across the plasma membrane. Potassium ion transport has been identified as a critical function for insect cold tolerance (MacMillan *et al*., 2016). These genes also showed signs of balancing selection (elevated nucleotide diversity and absolute differentiation), consistent with changes in gene function throughout development, generating opposing selection pressures that maintain higher levels of genetic variation in the population.

Together, our findings indicate that the evolution of thermoregulation in endotherms is shaped by life-stage-specific thermal vulnerabilities and demands. Understanding how animals will evolve and respond to climate change will therefore require increased focus on how thermal sensitivity and the molecular regulation of thermal tolerance shift across development.

## Methods

### Study site and animals

The experiment took place at the Western Cape Department of Agriculture’s ostrich research facility in Oudtshoorn, the Oudtshoorn Research Farm in the arid Klein Karoo of South Africa (GPS: 33°38′21.5′′S, 22°15′17.4′′E). Here, reproductive pairs of ostriches that are genetically similar to the Southern African subspecies (*Struthio camelus australis*) (Davids *et al*., 2012) and are commonly referred to as South African Blacks (*Struthio camelus* var. *domesticus*) are kept in enclosures of Karoo scrub with ad libitum water and feed. We obtained experimental ostrich chicks from adult pairs by collecting eggs within one day of laying and artificially incubating them at 36.2°C and a relative humidity of 24%. After hatching the chicks were reared in open enclosures (approximately 4 × 8 m) during the day and in stables at night. Chicks were fed with a predetermined plant-based pelleted diet consisting primarily of corn, soybean oilcake meal and alfalfa. In the three days preceding the experiment, all chicks were kept inside stables to ensure exposure to similar thermal environments.

### Experimental design and sample collection

To enable comparison of thermoregulatory responses across early development, we incubated two batches of eggs ∼7 weeks apart, such that 1-2 weeks old (“1-week-olds”, n = 18) and 7-9 weeks old (“8-week-olds”, n = 12) chicks of equal sex ratios could be simultaneously exposed to three experimental acute temperature treatments. The three temperature treatments were within the range of temperatures typically encountered in their natural habitat: cold 12°C), benign (23°C) and hot (40°C) (Fig 1D). Acute, controlled exposure of chicks to these temperatures was achieved by placing groups of chicks in three custom made climate chambers (approx. 3x4x2m) with temperatures controlled using air conditioner units (TCL portable air conditioner, 12000 BTU/h). On each experimental day, a mixed group consisting of six 1-week-olds (three males, three females) and four 8-week-olds (two males, two females) were placed in each of the three chambers at a starting temperature of ∼21°C. The temperature was then kept constant in one chamber to mimic a benign day, gradually raised over two hours in the second chamber to mimic a hot day, and gradually lowered over two hours in the third chamber to mimic a cold day (Fig 1D). After the treatment, at around 16:00, chicks were returned to the stable. This procedure was repeated on three consecutive days such that each group was exposed to all three temperature treatments in a randomised order, thereby minimising day effects, which were accounted for by including group as a random effect in statistical analyses. Four chicks died between experimental days and were replaced by chicks of the same age and sex (chick mortality is high during this age (Videvall *et al*., 2020)).

After three hours at the target temperatures, we collected blood samples for gene expression analysis within 30 seconds of removing the individuals from the chambers. We immediately transferred 50µl of blood into 500µl RNAlater and stored it first at 4°C and then at -20°C, to prevent RNA degradation (Harvey and Knutie, 2023). Gene expression from blood mRNA has been shown to be a good predictor of gene expression in other tissues (Basu *et al*., 2021) and is only mildly invasive, enabling multiple sampling of the same chicks over successive days. We measured cloacal temperature immediately after blood sampling to estimate the impact of the acute temperature treatment on their body temperature (Fig 1C). Chicks had access to food and water throughout the experiment.

### Cloacal temperature analysis

To test whether cloacal temperature was affected by the temperature treatments we fitted linear mixed models of cloacal temperature using lme4 (Bates *et al*., 2015) in R version 4.4.2 (R Core Team, 2024). Chicks that died during the experiment and their replacements were excluded from these analyses, resulting in 26 chicks. In the models, cloacal temperature was included as response variable, while fixed effects included sex (factorial: male or female) and temperature treatment (factorial: cold, benign or hot), both interacting with age (factorial: 1-week or 8-week), and random effects included chick ID and group (see Fig 1A) to account for differences in order of treatments and in chamber temperatures between treatment days. We then performed model comparisons against simpler models, first dropping the interactions, then sex, age and finally treatment, using the step function in lmerTest v 3.1-3 (Kuznetsova, Brockhoff and Christensen, 2017) which uses Satterthwaite’s method to identify the best fitting model. P-values were determined with a post hoc test using the Kenward-Roger method to estimate degrees of freedom, and adjusted with Tukey’s method using emmeans v 2.0.0 (Lenth *et al*., 2026) (see Table S1 for a summary of effect sizes).

To test if the individual cloacal temperature response to cold was related to the response to heat, we used a linear mixed model. The change in cloacal temperature from benign to cold was specified as the response variable; the change in cloacal temperature from benign to heat, age, sex and body mass (scaled to a mean of zero and a standard deviation of 1 within age group) were included as fixed effects, and group was included as a random effect. We then performed model comparisons against simpler models, first dropping sex, age, the change in cloacal temperature in heat, and body mass using the step function in lmerTest v 3.1-3 (Kuznetsova, Brockhoff and Christensen, 2017) that uses Satterthwaite’s method to identify the best fitting model. A p-value was obtained with an F-test using Satterthwaite’s method (see Table S2 for a summary of effect sizes). In all models we checked assumptions of parametric analyses using DHARMa v 0.4.7 and results were plotted using visreg v 2.8.0 (Breheny and Burchett, 2017) and ggplot v 4.0.0 (Wickham, 2016).

### RNA isolation, library preparation and sequencing

For sequencing, all samples from dead and replacement chicks were excluded, and samples from five chicks were excluded because of insufficient sample quality for library preparation. Consequently, transcriptomes were sequenced from 63 samples from 21 chicks – eleven 1-week-old (five males, six females) and ten 8-week-old chicks (five males, five females). RNA was extracted and isolated from 250µl 1:10 blood/RNAlater mix using the Qiagen RNeasy Mini Kit with Qiashredder (Hilden, Germany) according to protocol. To avoid contamination with DNA, samples were treated with Turbo DNase (Invitrogen, Carlsbad, CA). We assessed purity and concentration using a NanoDrop spectrophotometer (Thermo Scientific, USA). Additional quality control and sequencing was completed at Genewiz / Azenta Life Sciences, following their standard practice: RNA samples were quantified using a Qubit 4.0 Fluorometer (Life Technologies, Carlsbad, CA, USA) and RNA integrity was checked with an RNA Kit on Agilent 5300 Fragment Analyzer (Agilent Technologies, Palo Alto, CA, USA). mRNA sequencing libraries were prepared using the NEBNext Ultra II RNA Library Prep Kit for Illumina following manufacturer’s instructions (NEB, Ipswich, MA, USA). Sequencing libraries were validated using a NGS Kit on the Agilent 5300 Fragment Analyzer (Agilent Technologies, Palo Alto, CA, USA) and quantified by using a Qubit 4.0 Fluorometer (Invitrogen, Carlsbad, CA). The libraries were sequenced on Illumina NovaSeq 6000 using 2x150 Pair-End configuration v1.5. Raw sequence data (.bcl files) generated from Illumina NovaSeq was converted into fastq files and de-multiplexed using the Illumina bcl2fastq program version 2.20. One mismatch was allowed for index sequence identification.

### Genome annotation

There is limited ostrich RNA-Seq data available to guide annotation of the ostrich genome. We thus reannotated the *Struthio camelus australis* reference genome assembly downloaded from the DNA Zoo Consortium website (https://www.dnazoo.org, data from (Zhang *et al*., 2014)), combining RNA-Seq data from this study that was not used in differential gene expression analysis, and the European Nucleotide Archive using BRAKER 3.0.8 in ETP mode (study accessions PRJNA264356, PRJNA287589, PRJNA542010, PRJNA544815, PRJNA599522 and PRJNA646721. The vertebrate protein database OrthoDB was used as evidence for BRAKER (Kuznetsov *et al*., 2023). The quality of the annotation was assessed using BUSCO v 5.7.1 (Tegenfeldt *et al*., 2025) (Fig S1). Functional annotation was performed using eggNOG-mapper (version emapper-2.1.12) (Cantalapiedra *et al*., 2021) based on eggNOG orthology data (Huerta-Cepas *et al*., 2019). Sequence searches were performed using DIAMOND v 2.1.9 (Buchfink, Xie and Huson, 2015).

### Transcriptome analysis

Raw RNA-Seq reads were trimmed using the Trim Galore 0.6.10 (Krueger, 2023) wrapper tool, which uses Cutadapt 4.4 (Martin, 2011) for adaptor trimming and automatically checks read quality with FastQC 0.12.1 (Andrews, 2012). Read pairs with a Phred score under Q10 were discarded (Williams *et al*., 2016). Reads were mapped using STAR 2.7.11a (Dobin *et al*., 2013) to the *Struthio camelus australis* reference genome from the DNA Zoo Consortium website (https://www.dnazoo.org) using the new annotation. The number of reads uniquely mapping to features were counted using HTSeq 2.0.3 (Anders, Pyl and Huber, 2015; Putri *et al*., 2022). Globin and ferritin genes were removed because they had high counts that potentially reduce other signals. Because many reads mapped to globin and ferritin genes, coverage for 40 samples was lower than desired and they were thus re-sequenced using the same procedure as detailed above. Counts for each sample were summed across sequencing runs. This resulted in 3.5 M to 37 M uniquely mapped read counts per sample, with a median of 16M reads (excluding reads mapping to globin or ferritin). Three samples with outlier library sizes (37 M, 31 M and 3.5 M) were removed because they could bias dispersion estimation, resulting in 18 chicks with three samples, and three chicks with only two samples. As our analyses of gene expression focused on the change in expression within individuals across temperature treatments, we removed those three chicks missing one sample from further analysis, resulting in 18 chicks analysed (nine 1-week-olds and nine 8-week-olds).

We analysed differentially expressed genes using glmmSeq v0.5.5 (Lewis *et al*., 2022). Removing genes with low abundance across samples improves the detection of true differential expression (Bourgon, Gentleman and Huber, 2010), and we therefore filtered gene counts for genes with at least 10 reads in 10 samples. Size factors were calculated using trimmed mean of M-values normalisation using edgeR v 4.4.2 (Chen *et al*., 2024). Robust estimates of the negative binomial dispersion parameter were calculated for each gene (Zhou, Lindsay and Robinson, 2014). We checked for abnormalities and confounding factors using PCA analysis (Fig S2).

### Genetic basis of heat and cold responses

To identify genes that were differentially expressed between the treatments in the two age groups, we fitted generalised linear mixed effects models (GLMM) with a negative binomial distribution for each gene with temperature in the chambers as a categorical fixed effect (hot, benign and cold). We also included age (1 week or 8 week) and sex (male or female) as fixed effects. We included group (factorial: 1, 2 and 3) as random intercepts to account for differences in the order of exposure to the acute temperature treatments. Finally, we also included random slopes of chick ID across temperature (treatment | chick ID) to allow the impact of temperature treatment to vary between chicks. We removed 7 of the 11082 genes as their models failed to converge. Post hoc tests were performed on the genes significantly differentially expressed for temperature (q-value < 0.01) to determine whether genes were differentially expressed in heat vs benign or/and in cold vs benign in both age groups using the emmeans package v 2.0.0 (Lenth *et al*., 2026). Genes that were differentially expressed either in heat or cold (compared to benign) were categorised as showing a distinct response in heat and cold. Genes that were differentially expressed in both contrasts of heat vs benign and cold vs benign were categorised as having a shared response, i.e. they are involved both in the response to heat and to cold.

### Age modulation of heat and cold responses

To identify genes where differential expression in heat or cold were modulated by age, we constructed a second model that also included the interaction between age and temperature treatment. Hence the model had temperature treatment (hot, benign or cold), age (1-week or 8-week) and sex (male or female) as well as the age by temperature interaction as categorical fixed effects. The model also contained chick ID and group as random intercepts, and finally we also included random slopes of chick ID across temperature (treatment | chick ID). Post hoc tests were performed using the emmeans package v 2.0.0 (Lenth *et al*., 2026) on the genes significantly differentially expressed for the interaction term (q-value < 0.01), to identify if age modulated the gene expression response to heat, to cold or to both. Genes were grouped for heat and cold responses separately into being differentially expressed only in one age group (*only 1 week* or *only 8 week*), upregulated in one age group and downregulated in the other (*opposite direction*), or up- or downregulated in both age groups but with different intensities (*same direction*).

### Gene set over-representation analysis

Gene set over-representation analyses were carried out using clusterProfiler v 4.14.6 (Xu *et al*., 2024). Gene names from the functional gene annotation with eggNOG were matched to ENTREZ gene names in the *Gallus gallus* (chicken) database, leading to a gene universe of 10471 genes. Then, enriched KEGG and GO terms in different gene sets with an adjusted p-value < 0.1 were identified using enrichKEGG and enrichGO functions. Differentially expressed genes from the models with and without age-by-treatment interactions were analysed separately to highlight differences between genes modulated by age and those not modulated by age.

### Detecting signatures of selection using whole genome sequencing

To test if genes involved in the response to heat and cold exposure have been important for local adaptation in natural populations, we analysed individual whole genome sequences of 20 individuals from populations of two subspecies: A population of the Masai ostrich (*S. c. massaicus*), also referred to as Kenyan Red, established at the research site in 2007 (n_founders_ = 18), and the South African ostrich (*S. c. australis*), also referred to as the Zimbabwean Blues because of its origin in Namibia and Zimbabwe, established in 1990 (n_founders_ = 83). To aid variant calling and population structure analyses we included already published genetic data from 10 individuals of the Southern Africa Blacks (*S. c.* var. *domesticus*) (Yazdi *et al*., 2023) in the analysis, a third population at the research site established in 1990 (n_founders_ = 171), which was used for the gene expression experiment. These samples were however not used for population differentiation analyses because of likely introgression events of Zimbabwean Blues into Southern Africa Blacks (Davids *et al*., 2012). All samples were sequenced at the Science for Life Laboratory, National Genomics Infrastructure, using Illumina HiSeq 2500, following the manufacturer’s protocol.

### Population differentiation analyses

Reads were trimmed and adapters removed using Trim Galore v 0.6.10. Reads with a Phred score < 20 were discarded and Ns clipped from sequences. Then reads were aligned to the *Struthio camelus australis* reference genome from the DNA Zoo Consortium website (https://www.dnazoo.org) with the new annotation (see supplementary information for read statistics) using bwa-mem v 0.7.18 (Li and Durbin, 2009) and samtools v 1.22 was used to remove duplicates and filter for reads with a MQ > 20 (Danecek *et al*., 2021). Scaffolds < 1000bp were discarded from further analysis (accounting to 0.16% of the genome). Variants were called using freebayes v 1.3.6 in 1MB segments (Garrison and Marth, 2012). The -- populations option was used to inform freebayes of subspecies membership. The population-based Bayesian inference model is thus partitioned based on the populations. Finally, the vcf files of the 1 MB genome chunks were concatenated into one file for statistics and filtering.

The sites used for subspecies differentiation were filtered using vcftools v 0.1.17 (Danecek *et al*., 2011) and bcftools v 1.22 (Danecek *et al*., 2021). Sites were filtered for a mean depth of at least a third of average depth (10X) and at most twice average depth (60X), and sites missing in more than 10% of samples (i.e. three individuals) were removed. Indels and multiallelic sites removed next. Sites out of Hardy-Weinberg equilibrium (HWE) with p < 0.01 were determined separately for each subspecies, and analyses were performed both with and without removing these sites, which led to very similar results; only results without removing sites out of HWE are shown.

To validate the subspecies assignment of each individual we performed population structure analysis. For this we removed monomorphic sites, sites with QUAL below 20 and minor allele frequency below 3% (i.e. singletons). We linkage pruned SNPs using Plink v 1.90b7.7 in 50 kb windows with a 10 bp step size, allowing a correlation of r^2^ < 0.1 (Purcell *et al*., 2007). We constructed a PCA of the individual samples using Plink (Fig S4). We also ran ADMIXTURE v 1.3.0 for k=2,…,5 (Alexander, Novembre and Lange, 2009) (Fig S5). Based on the results we removed two individuals – one South African ostrich and one Masai ostrich – due to unclear population assignment. Then we performed relatedness analysis (Fig S6) and found four individuals that were closely related to others (full sibs and parents) and removed these as well (one South African ostrich and three Masai ostriches) resulting in ten Southern Africa Blacks, eight South African ostriches and six Masai ostriches available for analysis.

To test for differentiation that could stem from different thermal environments, we compared South African ostriches and Masai ostriches. Hudson F_ST_, d_xy_ and π were determined in the genes differentially expressed in response to heat and cold, and in the genes whose response to heat and cold was modulated by age, using scripts by Simon Martin (https://github.com/simonhmartin/genomics_general, accessed on 16.09.2025). Each gene was expanded with 500bp upstream of the start codon to capture genetic variation in the promoter region. Other promoter sizes (100-1000bp) were also tested, with similar results. Only regions on scaffolds > 1000 bp were analysed, which excluded 5 regions. Regions with < 10 SNPs were discarded from these analyses, which excluded 38 genes. We tested for differences in Hudson F_ST_ and d_xy_ between groups of genes using linear models with gene group as a fixed effect. We tested for differences in pi between groups of genes using linear mixed models with gene group and population as fixed effects and gene ID as random effect, via REML using lme4 v 1.1-37 (Bates *et al*., 2015). Subsequently, we determined least-square means and pairwise contrasts using the Kenward-Roger degrees of freedom method and Tukey’s p-value adjustment using emmeans v 2.0.0 (Lenth *et al*., 2026) in R. Tests among gene sets that were up- or downregulated in heat and/or cold included 253 genes, while tests among gene sets with different types of age-modulation included 373 genes.

### Ethics

All procedures were approved by the Departmental Ethics Committee for Research on Animals (DECRA) of the Western Cape Department of Agriculture, reference no. AP/NP/O/AE117.

## Acknowledgements

We thank the staff and workers at the Oudtshoorn Research Farm for their assistance with data collection and maintenance of the ostriches, and we thank the Western Cape Government for use of their resources. We thank Marie Rosenstand Hansen for her assistance with the laboratory work related to RNA extraction.

## Author contributions

Conceptualisation: M.F.S., C.K.C.; Data curation: M.F.S., L.L.H., Z.B., S.C., A.E.; Formal analysis: L.L.H., M.F.S., Funding acquisition: M.F.S.; Investigation: M.F.S., L.L.H., M.K., C.K.C., N.L., A.E.; Methodology: L.L.H., M.F.S., C.K.C.; Project administration: M.F.S.; Writing-original draft: L.L.H., M.F.S., C.K.C. Writing –reviewing and editing: L.L.H., M.F.S., C.K.C., M.K., N.L., A.E., Z.B., S.C.

## Data availability

The genetic data that support the findings of this study are available on the NCBI BioSample database under accessions SAMN55035168 to SAMN55035230 and SAMN56364120 to SAMN56364139; the morphological data can be found on Figshare under https://doi.org/10.6084/m9.figshare.31424045.

## Code availability

Code for the analysis is available on GitHub at https://github.com/lealhuber under the repositories ChamberTempRNA, Subspecies, ostrich-annotation and DGE_Rcode.

## References

Alexander, D.H., Novembre, J. and Lange, K. (2009) “Fast model-based estimation of ancestry in unrelated individuals,” Genome Research, 19(9), pp. 1655–1664. Available at: 10.1101/gr.094052.109.

Anders, S., Pyl, P.T. and Huber, W. (2015) “HTSeq—a Python framework to work with high-throughput sequencing data,” Bioinformatics, 31(2), pp. 166–169. Available at: 10.1093/bioinformatics/btu638.

Andrews, S. (2012) “FastQC: a quality control tool for high throughput sequence data.” Available at: http://www.bioinformatics.babraham.ac.uk/projects/fastqc.

Ballinger, M.A. et al. (2023) “Environmentally robust cis-regulatory changes underlie rapid climatic adaptation,” Proceedings of the National Academy of Sciences, 120(39), p. e2214614120. Available at: 10.1073/pnas.2214614120.

Basu, M. et al. (2021) “Predicting tissue-specific gene expression from whole blood transcriptome,” Science Advances, 7(14). Available at: 10.1126/sciadv.abd6991.

Bates, D. et al. (2015) “Fitting Linear Mixed-Effects Models Using lme4,” Journal of Statistical Software, 67(1). Available at: 10.18637/jss.v067.i01.

Bathiany, S. et al. (2018) “Climate models predict increasing temperature variability in poor countries,” Science Advances, 4(5), p. eaar5809. Available at: 10.1126/sciadv.aar5809.

Bourgon, R., Gentleman, R. and Huber, W. (2010) “Independent filtering increases detection power for high-throughput experiments,” Proceedings of the National Academy of Sciences, 107(21), pp. 9546–9551. Available at: 10.1073/pnas.0914005107.

Breheny, P. and Burchett, W. (2017) “Visualization of Regression Models Using visreg,” The R Journal, 9(2), pp. 56–71. Available at: 10.32614/RJ-2017-046.

Buchfink, B., Xie, C. and Huson, D.H. (2015) “Fast and sensitive protein alignment using DIAMOND,” Nature Methods, 12(1), pp. 59–60. Available at: 10.1038/nmeth.3176.

Cantalapiedra, C.P., et al. (2021) “eggNOG-mapper v2: Functional Annotation, Orthology Assignments, and Domain Prediction at the Metagenomic Scale,” Molecular Biology and Evolution. Edited by K. Tamura, 38(12), pp. 5825–5829. Available at: 10.1093/molbev/msab293.

Chen, B. et al. (2017) “Transposable Element-Mediated Balancing Selection at Hsp90 Underlies Embryo Developmental Variation,” Molecular Biology and Evolution, 34(5), pp. 1127–1139. Available at: 10.1093/molbev/msx062.

Chen, P. and Zhang, J. (2020) “Antagonistic pleiotropy conceals molecular adaptations in changing environments,” Nature Ecology & Evolution, 4(3), pp. 461–469. Available at: 10.1038/s41559-020-1107-8.

Chen, Y. et al. (2024) “edgeR v4: powerful differential analysis of sequencing data with expanded functionality and improved support for small counts and larger datasets.” bioRxiv, p. 2024.01.21.576131. Available at: 10.1101/2024.01.21.576131.

Craig, E.A. and Schlesinger, M.J. (1985) “The Heat Shock Response,” Critical Reviews in Biochemistry, 18(3), pp. 239–280. Available at: 10.3109/10409238509085135.

Cruickshank, T.E. and Hahn, M.W. (2014) “Reanalysis suggests that genomic islands of speciation are due to reduced diversity, not reduced gene flow,” Molecular Ecology, 23(13), pp. 3133–3157. Available at: 10.1111/mec.12796.

Dahlke, F.T. et al. (2020) “Thermal bottlenecks in the life cycle define climate vulnerability of fish,” Science, 369(6499), pp. 65–70. Available at: 10.1126/science.aaz3658.

Danecek, P. et al. (2011) “The variant call format and VCFtools,” Bioinformatics, 27(15), pp. 2156–2158. Available at: 10.1093/bioinformatics/btr330.

Danecek, P. et al. (2021) “Twelve years of SAMtools and BCFtools,” GigaScience, 10(2), p. giab008. Available at: 10.1093/gigascience/giab008.

Davids, A. et al. (2012) “Genetic variation within and among three ostrich breeds, estimated by using microsatellite markers,” South African Journal of Animal Science, 42(2), pp. 156–163. Available at: 10.4314/sajas.v42i2.8.

Dobin, A. et al. (2013) “STAR: ultrafast universal RNA-seq aligner,” Bioinformatics, 29(1), pp. 15–21. Available at: 10.1093/bioinformatics/bts635.

Fick, S.E. and Hijmans, R.J. (2017) “WorldClim 2: new 1-km spatial resolution climate surfaces for global land areas,” International Journal of Climatology, 37(12), pp. 4302–4315. Available at: 10.1002/joc.5086.

Fischer, E.M., Sippel, S. and Knutti, R. (2021) “Increasing probability of record-shattering climate extremes,” Nature Climate Change, 11(8), pp. 689–695. Available at: 10.1038/s41558-021-01092-9.

Garrison, E. and Marth, G. (2012) “Haplotype-based variant detection from short-read sequencing.” arXiv. Available at: 10.48550/arXiv.1207.3907.

Gething, M.-J. and Sambrook, J. (1992) “Protein folding in the cell,” Nature, 355(6355), pp. 33–45. Available at: 10.1038/355033a0.

Harvey, J.A. and Knutie, S.A. (2023) “Effect of RNA preservation methods on RNA quantity and quality of field-collected avian whole blood,” Avian Biology Research, 16(2), pp. 51–58. Available at: 10.1177/17581559231169179.

Huerta-Cepas, J. et al. (2019) “eggNOG 5.0: a hierarchical, functionally and phylogenetically annotated orthology resource based on 5090 organisms and 2502 viruses,” Nucleic Acids Research, 47(D1), pp. D309–D314. Available at: 10.1093/nar/gky1085.

Junprung, W. et al. (2021) “Balancing selection at the ATP binding site of heat shock cognate 70 (HSC70) contributes to increased thermotolerance in *Artemia franciscana*,” Aquaculture, 531, p. 735988. Available at: 10.1016/j.aquaculture.2020.735988.

Kawecki, T.J. et al. (2021) “The Genomic Architecture of Adaptation to Larval Malnutrition Points to a Trade-off with Adult Starvation Resistance in Drosophila,” Molecular Biology and Evolution, 38(7), pp. 2732–2749. Available at: 10.1093/molbev/msab061.

Kim, Hana et al. (2021) “Transcriptomic Response under Heat Stress in Chickens Revealed the Regulation of Genes and Alteration of Metabolism to Maintain Homeostasis,” Animals, 11(8), p. 2241. Available at: 10.3390/ani11082241.

Krueger, F. (2023) “Trim Galore!” Available at: 10.5281/zenodo.5127898.

Kuznetsov, D. et al. (2023) “OrthoDB v11: annotation of orthologs in the widest sampling of organismal diversity,” Nucleic Acids Research, 51(D1), pp. D445–D451. Available at: 10.1093/nar/gkac998.

Kuznetsova, A., Brockhoff, P.B. and Christensen, R.H.B. (2017) “lmerTest Package: Tests in Linear Mixed Effects Models,” Journal of Statistical Software, 82(13). Available at: 10.18637/jss.v082.i13.

Lenth, R.V. et al. (2026) “emmeans: Estimated Marginal Means, aka Least-Squares Means.” Available at: https://cran.r-project.org/web/packages/emmeans/index.html (Accessed: April 27, 2026).

Lewis, M. et al. (2022) “glmmSeq: General Linear Mixed Models for Gene-Level Differential Expression.” Available at: https://myles-lewis.github.io/glmmSeq/.

Li, H. and Durbin, R. (2009) “Fast and accurate short read alignment with Burrows–Wheeler transform,” Bioinformatics, 25(14), pp. 1754–1760. Available at: 10.1093/bioinformatics/btp324.

Liu, Q. et al. (2025) “Molecular Pathogenesis of Avian Splenic Injury Under Thermal Challenge: Integrated Mitigation Strategies for Poultry Heat Stress,” Current Issues in Molecular Biology, 47(6), p. 410. Available at: 10.3390/cimb47060410.

MacMillan, H.A. et al. (2016) “Preservation of potassium balance is strongly associated with insect cold tolerance in the field: a seasonal study of Drosophila subobscura,” Biology Letters, 12(5), p. 20160123. Available at: 10.1098/rsbl.2016.0123.

Martin, M. (2011) “Cutadapt removes adapter sequences from high-throughput sequencing reads,” EMBnet.journal, 17.1, pp. 10–12. Available at: 10.14806/ej.17.1.200.

Monson, M.S. et al. (2018) “Immunomodulatory effects of heat stress and lipopolysaccharide on the bursal transcriptome in two distinct chicken lines,” BMC Genomics, 19(1), p. 643. Available at: 10.1186/s12864-018-5033-y.

Perkins-Kirkpatrick, S.E. and Lewis, S.C. (2020) “Increasing trends in regional heatwaves,” Nature Communications, 11(1), p. 3357. Available at: 10.1038/s41467-020-16970-7.

Purcell, S. et al. (2007) “PLINK: a tool set for whole-genome association and population-based linkage analyses,” American Journal of Human Genetics, 81(3), pp. 559–575. Available at: 10.1086/519795.

Putri, G.H., et al. (2022) “Analysing high-throughput sequencing data in Python with HTSeq 2.0,” Bioinformatics. Edited by V. Boeva, 38(10), pp. 2943–2945. Available at: 10.1093/bioinformatics/btac166.

R Core Team (2024) “R: A Language and Environment for Statistical Computing.” Vienna, Austria: R Foundation for Statistical Computing. Available at: https://www.R-project.org/.

Riddell, E.A. et al. (2019) “Cooling requirements fueled the collapse of a desert bird community from climate change,” Proceedings of the National Academy of Sciences, 116(43), pp. 21609–21615. Available at: 10.1073/pnas.1908791116.

Rodríguez-Verdugo, A. et al. (2014) “Different tradeoffs result from alternate genetic adaptations to a common environment,” Proceedings of the National Academy of Sciences, 111(33), pp. 12121–12126. Available at: 10.1073/pnas.1406886111.

Rotenberry, J.T. and Balasubramaniam, P. (2020) “Estimating egg mass–body mass relationships in birds,” The Auk, 137(3), pp. 1–11. Available at: 10.1093/auk/ukaa019.

Rowe, M.F. et al. (2013) “Heat storage in Asian elephants during submaximal exercise: behavioral regulation of thermoregulatory constraints on activity in endothermic gigantotherms,” Journal of Experimental Biology, 216(10), pp. 1774–1785. Available at: 10.1242/jeb.076521.

Scholander, P.F. et al. (1950) “Heat regulation in some arctic and tropical mammals and birds,” The Biological Bulletin, 99(2), pp. 237–258. Available at: 10.2307/1538741.

Schou, M.F. et al. (2022) “Evolutionary trade-offs between heat and cold tolerance limit responses to fluctuating climates,” Science Advances, 8(21). Available at: 10.1126/sciadv.abn9580.

Schou, M.F. and Cornwallis, C.K. (2024) “Adaptation to fluctuating temperatures across life stages in endotherms,” Trends in Ecology & Evolution, 39(9), pp. 841–850. Available at: 10.1016/j.tree.2024.05.012.

Sørensen, J.G. et al. (2016) “Thermal fluctuations affect the transcriptome through mechanisms independent of average temperature,” Scientific Reports, 6(1), p. 30975. Available at: 10.1038/srep30975.

Svensson, E.I. et al. (2023) “Heritable variation in thermal profiles is associated with reproductive success in the world’s largest bird,” Evolution Letters [Preprint]. Available at: 10.1093/evlett/qrad049.

Tang, L.-P. et al. (2021) “Heat stress inhibits expression of the cytokines, and NF-κB-NLRP3 signaling pathway in broiler chickens infected with *salmonella typhimurium*,” Journal of Thermal Biology, 98, p. 102945. Available at: 10.1016/j.jtherbio.2021.102945.

Tegenfeldt, F. et al. (2025) “OrthoDB and BUSCO update: annotation of orthologs with wider sampling of genomes,” Nucleic Acids Research, 53(D1), pp. D516–D522. Available at: 10.1093/nar/gkae987.

Videvall, E. et al. (2020) “Early-life gut dysbiosis linked to juvenile mortality in ostriches,” Microbiome, 8(1), p. 147. Available at: 10.1186/s40168-020-00925-7.

Wickham, H. (2016) ggplot2: Elegant Graphics for Data Analysis. New York: Springer-Verlag. Available at: https://ggplot2.tidyverse.org.

Williams, C.R. et al. (2016) “Trimming of sequence reads alters RNA-Seq gene expression estimates,” BMC Bioinformatics, 17(1), p. 103. Available at: 10.1186/s12859-016-0956-2.

Xie, S., Tearle, R. and McWhorter, T.J. (2018) “Heat Shock Protein Expression is Upregulated after Acute Heat Exposure in Three Species of Australian Desert Birds,” Avian Biology Research, 11(4), pp. 263–273. Available at: 10.3184/175815618X15366607700458.

Xu, S. et al. (2024) “Using clusterProfiler to characterize multiomics data,” Nature Protocols, 19(11), pp. 3292–3320. Available at: 10.1038/s41596-024-01020-z.

Yazdi, H.P., et al. (2023) “The evolutionary maintenance of ancient recombining sex chromosomes in the ostrich,” PLOS Genetics. Edited by G. Bosco, 19(6), p. e1010801. Available at: 10.1371/journal.pgen.1010801.

Yin, C. et al. (2025) “Regulation of gene expression under temperature stress and genome-wide analysis of heat shock protein family in *Eriocheir sinensis*,” International Journal of Biological Macromolecules, 308, p. 142503. Available at: 10.1016/j.ijbiomac.2025.142503.

Zhang, G. et al. (2014) “Comparative genomics reveals insights into avian genome evolution and adaptation,” Science, 346(6215), pp. 1311–1320. Available at: 10.1126/science.1251385.

Zhou, X., Lindsay, H. and Robinson, M.D. (2014) “Robustly detecting differential expression in RNA sequencing data using observation weights,” Nucleic Acids Research, 42(11), p. e91. Available at: 10.1093/nar/gku310.

