## Supplemental information for "Development shapes molecular responses to thermal extremes in a desert bird"

### Supplementary Information

Chamber Temperature RNA Experiment

07 August, 2026

### 1 Figures

#### 1.1 Figure S1: BUSCO analysis of *Struthio camelus* genome annotation

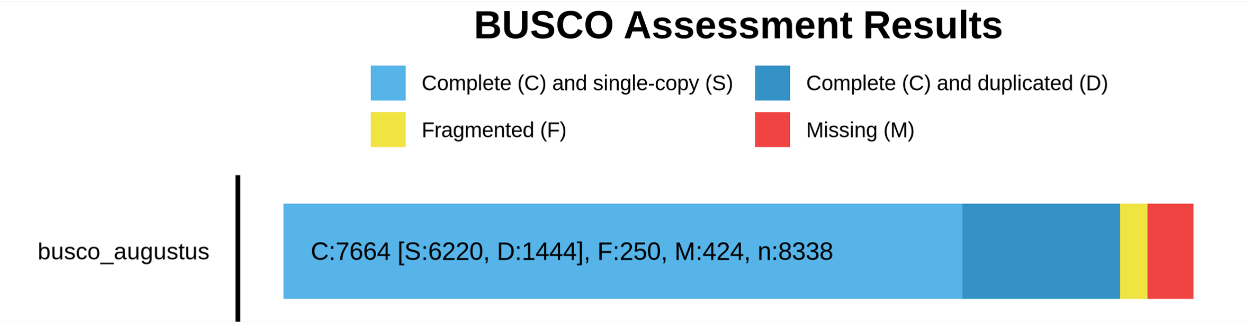

#### 1.2 Figure S2: PCA of differentially expressed genes

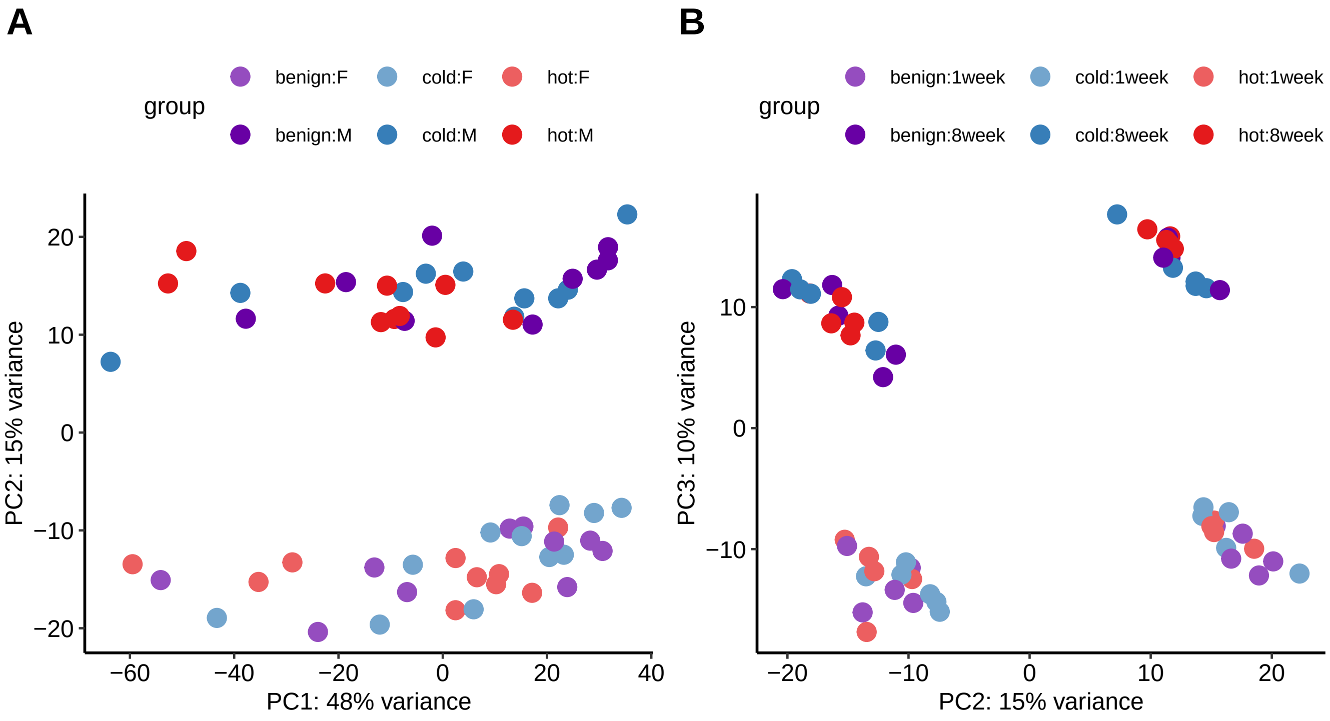

PCA of normalised and variance stabilised gene counts of only the used samples, using the 1000 most variable genes. A) PC1 and PC2, B) PC2 and PC3. M refers to males, F to females. PC2 separates the sexes, and PC3 separates the age groups.

##### 1.3 Figure S3: Expression in heat and cold

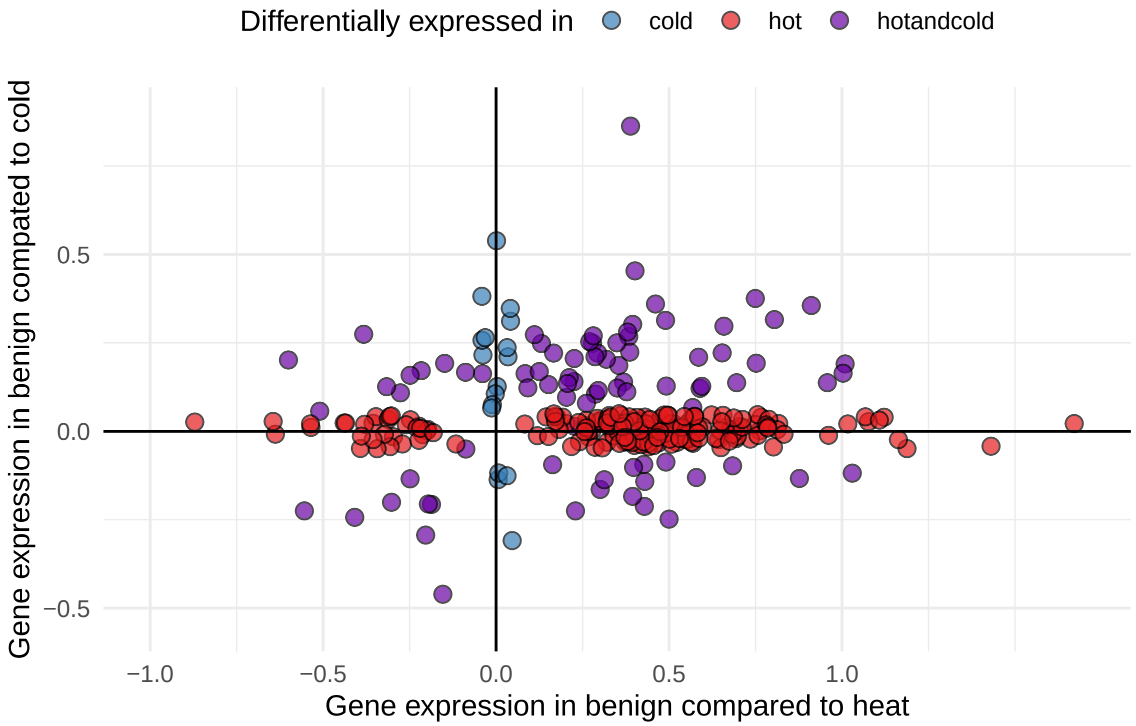

Plot of modelled gene expression in benign - cold and benign - heat, coloured by significance. Only genes with no age interaction are shown. The non-significant effects in cold of the genes only differentially expressed in heat, and in heat of the genes only DE in cold, were set to 0 and then jittered by 0.05.

#### 1.4 Figure S4: PCA plot of *S.camelus* subspecies

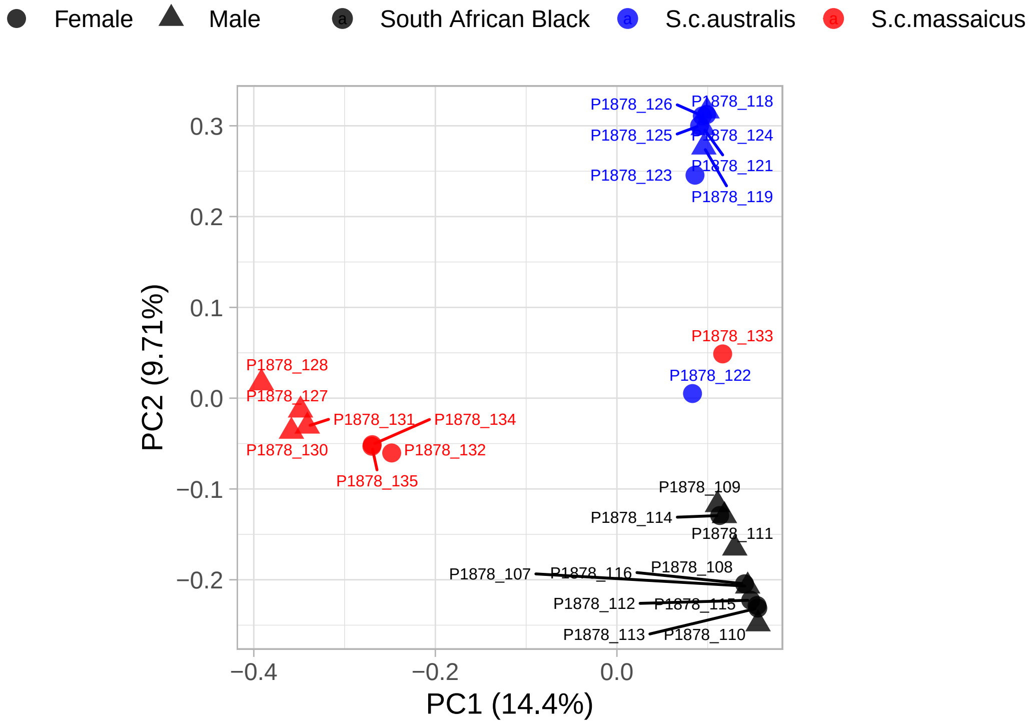

PCA based on whole genome sequencing data. PC1 separates *S.c. massaicus* from the other two populations, which are separated by PC2. Individuals P1878_122 and P1878_133 seem to be mixes of *S.c. australis* and the farmed "South African Black" breed and did not correspond with their labelling and were thus removed from further analysis.

#### 1.5 Figure S5: Admixture plot of *S.camelus* subspecies

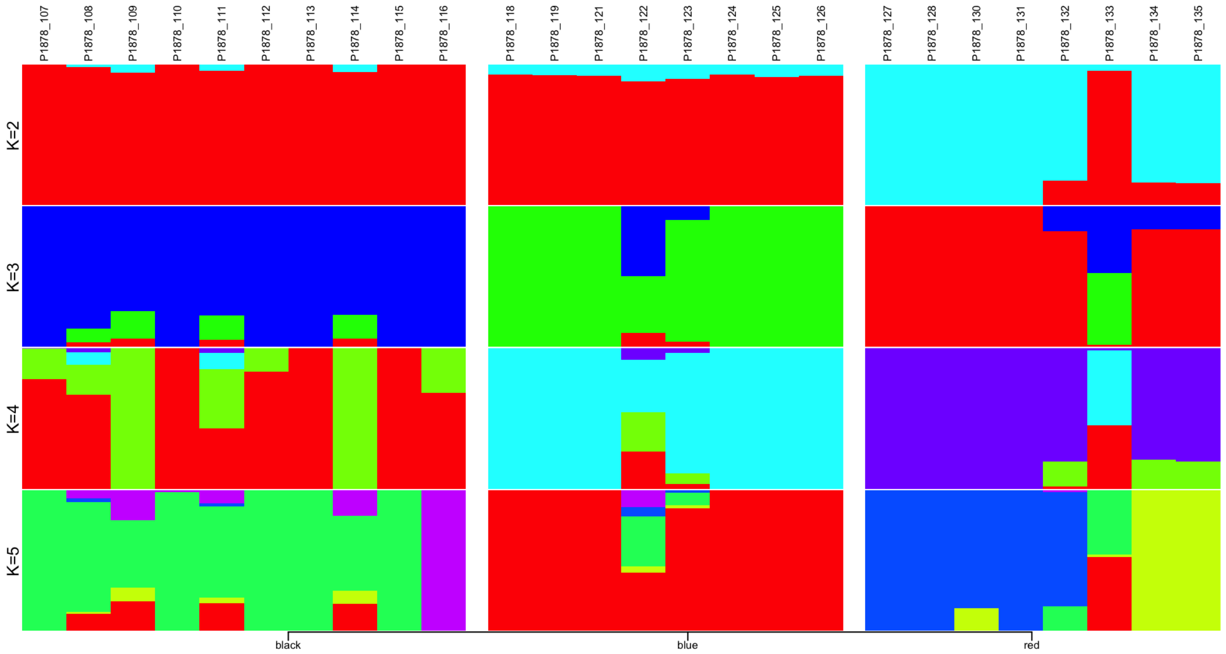

The ADMIXTURE results reflect the PCA results very well, showing first the separation of *S.c. massaicus* and at k=2 and the other two at k=3, and supports that P1878_122 and P1878_133 are mixes.

#### 1.6 Figure S6: Realatedness analysis

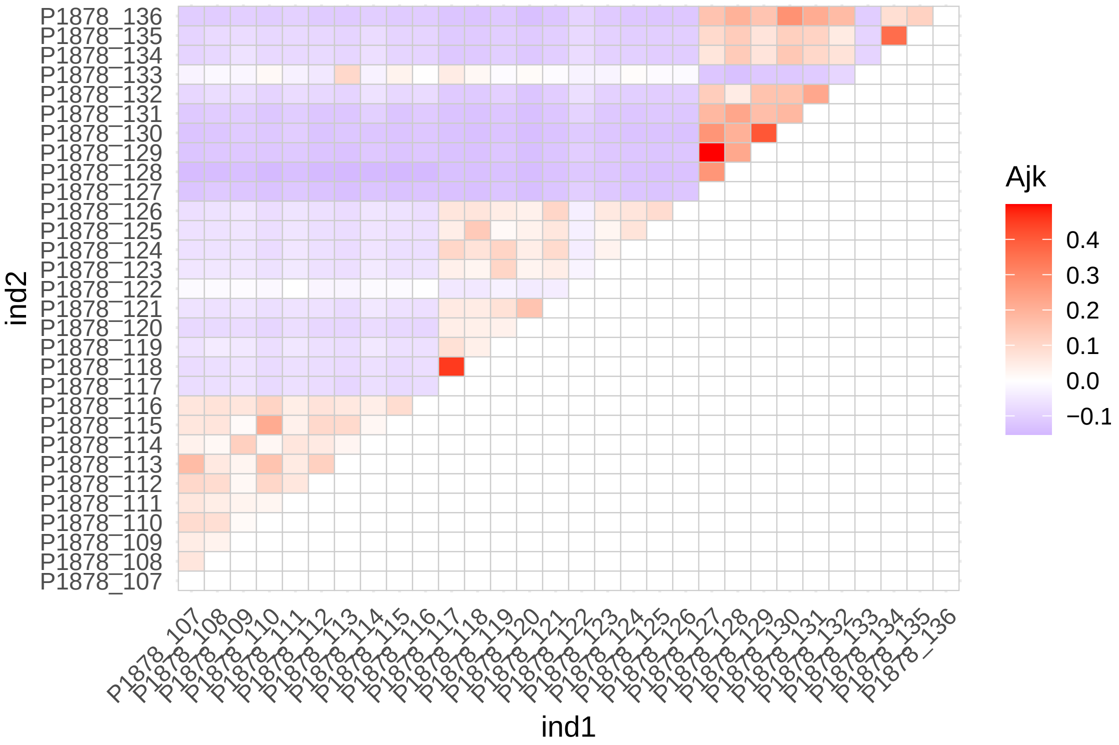

After depth, missingness and indel filtering, vcftools –relatedness was run which calculates a relatedness statistic based on the method of Yang et al, Nature Genetics 2010 (doi:10.1038/ng.608), called the unadjusted Ajk statistic. Expectation of Ajk is zero for individuals within a population, and one for an individual with themselves (but the diagonal was set to 0 here to make the colours better distinguishable). Based on this, individuals P1878_117, P1878_129, P1878_134 and P1878_136 were removed from further analysis.

##### 1.7 Figure S7: Pi and Dxy between *S.c.australis* and *S.c.massaicus* in thermoregulatory genes

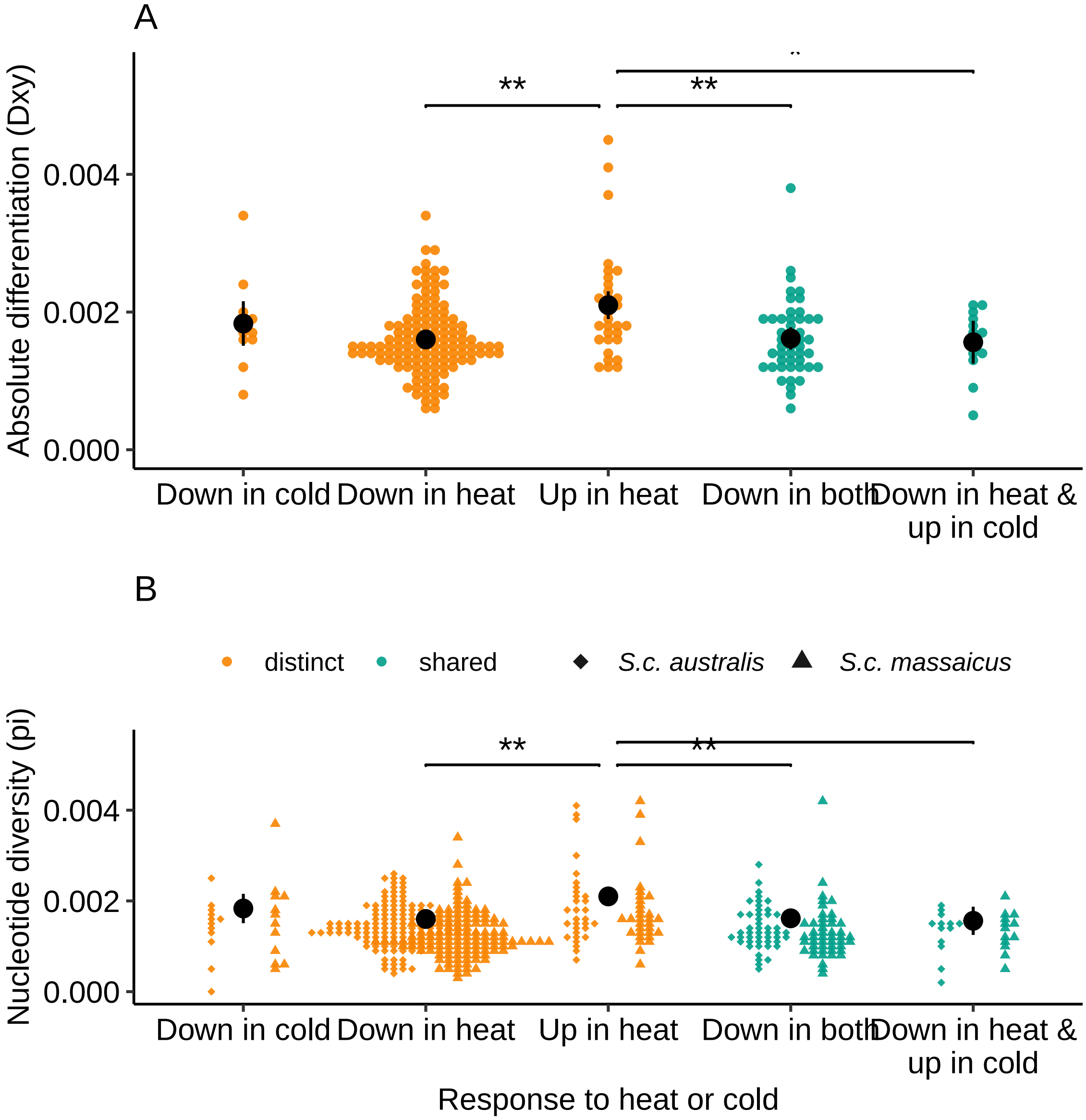

*Genes involved in thermoregulation carry signals of balancing selection across ostrich subspecies*: Nucleotide diversity (pi) and absolute differentiation (Dxy) of the South African and Masai subspecies are higher in genes that respond to the same temperature across ages but are upregulated in one age and downregulated in the other, as well as genes upregulated in response to heat. Significant differences between categories were determined using a linear mixed model and emmeans. Only gene sets with more than 10 genes and only genes with $> 10$ SNPs were included in the analyses.

(**A-B**): Pi and Dxy of genes only responding to either heat or cold exposure in both age groups (distinct) and genes responding to both heat and cold (shared). Distinct genes are grouped into genes downregulated in cold compared to benign (colddown) and genes downregulated (hotdown) or upregulated (hotup) in heat. Shared genes are grouped into genes downregulated both in heat and cold compared to benign (bothdown), and genes downregulated in heat but upregulated in cold (hotdowncoldup) (see also Fig 2B).

#### 1.8 Figure S8: Fst between *S.c.australis* and *S.c.massaicus* in thermoregulatory genes

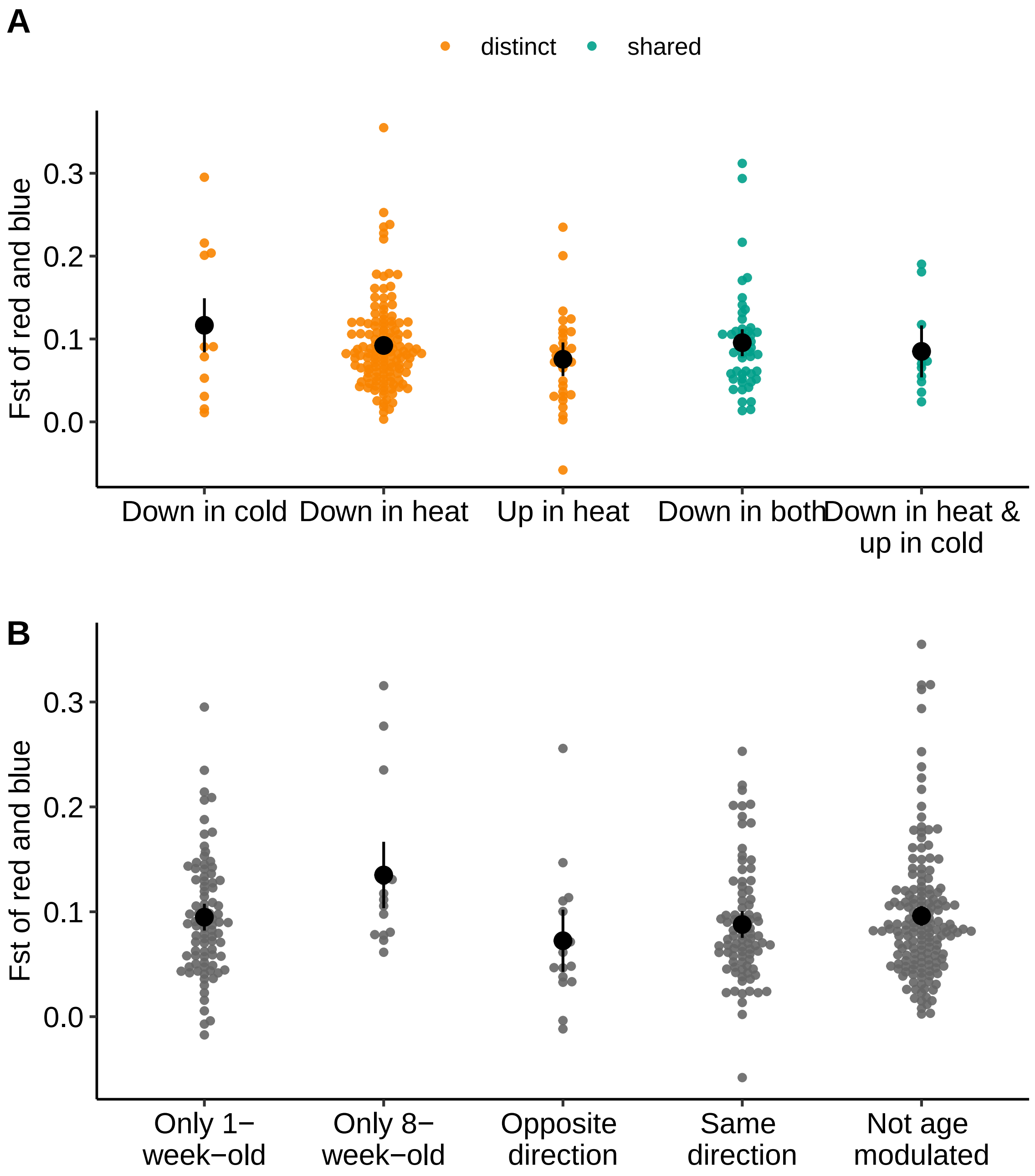

A) Genes categorised by being shared or distinct and up or downregulated in heat and cold.

B) Genes categorised by being differentially expressed in heat in one age group or both in opposite or the same direction. Only categories with > 10 genes are shown, and only genes with > 10 SNPs. There were no significant contrasts. Note that $F_{ST}$ results can be biased by differences in nucleotide diversity between populations, which is the case here as *S.c.massaicus* has consistently lower $\pi$ than *S.c.australis*, likely because they stem from a small founding population.

#### 1.9 Figure S9: Influence of gene length on pi and dxy

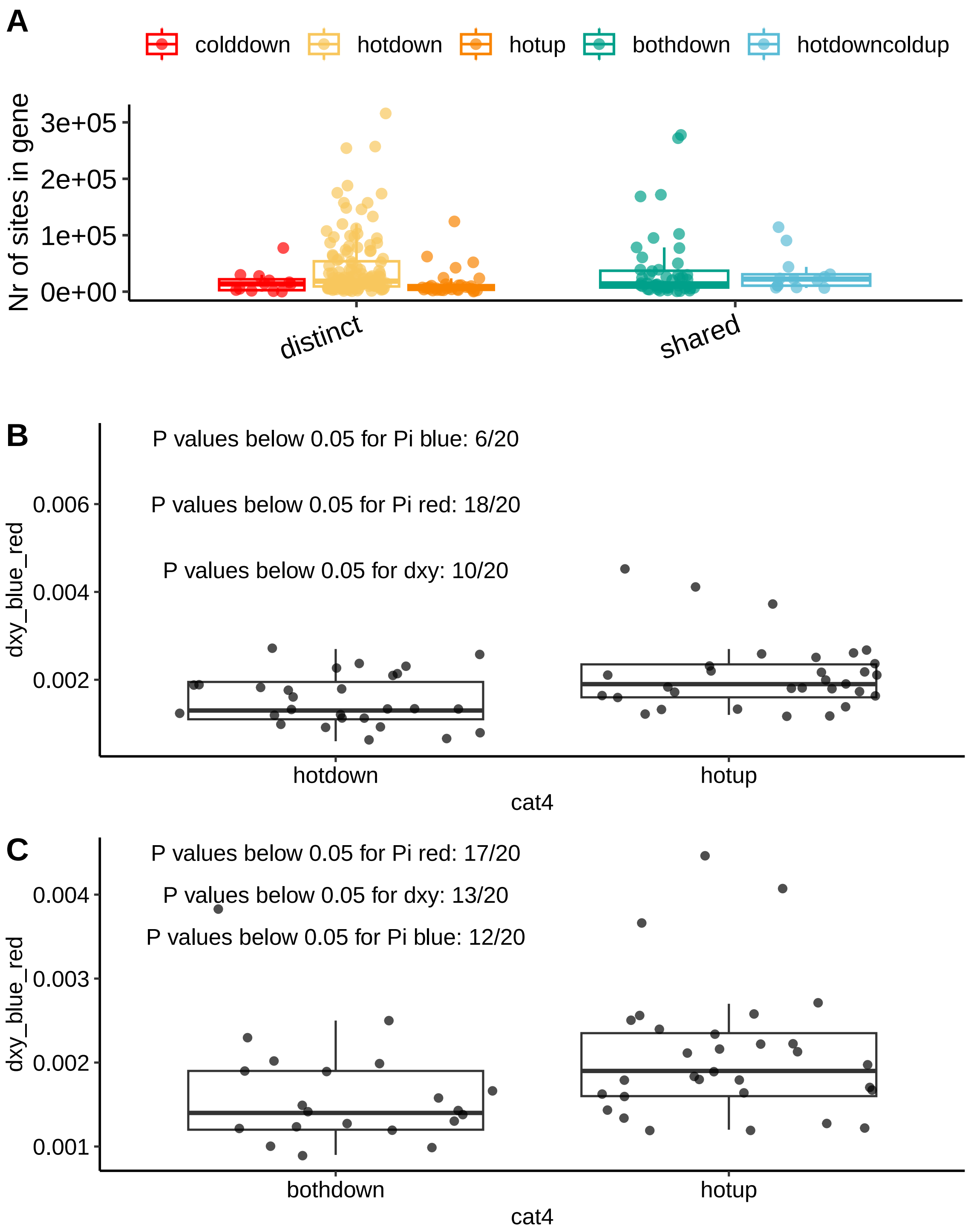

Gene length or the number of SNPs per gene could influence the $\pi$, $d_{xy}$ or $F_{ST}$ results, so we are checking here if it does. The results indicate that differences in gene lengths do not influence the results.

**A)** number of sites per gene in temperature related genes

**B)** Brunnermunzel p-values of pi and dxy between hotdown and hotup after sampling a similar distribution of number of sites from hotdown as hotup

**C)** Brunnermunzel p-values of pi and dxy between bothdown and hotup after sampling a similar distribution of number of sites from bothdown as hotup

### 2 Tables

#### 2.1 Table S1: Cloacal temperature model

Table 1: Linear mixed model fit by REML. F-tests use Satterthwaite's method. Formula: cloacat treatment*age + treatment*sex + (1|chickno) + (1|group). REML = 105.2

| **Parameter** | **Eliminated** | **Df** | **Sum of Sq** | **RSS** | **AIC** | **F value** | **Pr(>F)** |
|---|---|---|---|---|---|---|---|
| treatment:sex | 1 | 2 | 0.114 | 20.553 | -90.029 | 0.192 | 0.826 |
| treatment:Age | 2 | 2 | 0.292 | 20.845 | -92.928 | 0.505 | 0.606 |
| sex | 3 | 1 | 0.115 | 20.961 | -94.497 | 0.404 | 0.527 |
| Age | 4 | 1 | 0.483 | 21.443 | -94.722 | 1.703 | 0.196 |
| treatment | 0 | 2 | 53.564 | 75.007 | -1.052 | 93.674 | 0 |
| **Type** | **Parameter** | **Estimate** | **Std.Err/Dev** | **df** | | | |
| Fixed effects | (Intercept) | 39.286 | 0.133 | 4.421 | | | |
| Fixed effects | treatmentcold | -0.192 | 0.138 | 50 | | | |
| Fixed effects | treatmenthot | 1.654 | 0.138 | 50 | | | |
| Fixed effects | marginal R squared | 0.701 | | | | | |
| Random effect var. | chickno | 0.025 | 0.158 | | | | |
| Random effect var. | group | 0.022 | 0.148 | | | | |
| Random effect var. | Residual | 0.247 | 0.497 | | | | |
| **contrast** | **estimate** | **SE** | **df** | **t.ratio** | **pvalue** | | |
| benign - cold | 0.192 | 0.138 | 50 | 1.396 | 0.3509 | | |
| benign - hot | -1.654 | 0.138 | 50 | -12.002 | 0 | | |
| cold - hot | -1.846 | 0.138 | 50 | -13.397 | 0 | | |

#### 2.2 Table S2: Model of change in cloacal temperature

Table 2: Linear mixed model fit by REML. F-tests use Satterthwaite's method. Formula: TchangeCold TchangeHot + (1|group). REML = 37.4

| **Parameter** | **Eliminated** | **Sum Sq** | **Mean Sq** | **NumDF** | **DenDF** | **F value** | **Pr(>F)** |
|---|---|---|---|---|---|---|---|
| sex | 1 | 0.006 | 0.006 | 1 | 19.233 | 0.024 | 0.878 |
| Age | 2 | 0.118 | 0.118 | 1 | 20.2 | 0.476 | 0.498 |
| TchangeHot | 0 | 2.308 | 2.308 | 1 | 22.977 | 9.515 | 0.005 |
| massScaled | 0 | 1.049 | 1.049 | 1 | 21.327 | 4.323 | 0.05 |
| **Type** | **Estimate** | **Std. Error** | **df** | | | | |
| (Intercept) | -0.955 | 0.332 | 8.203 | | | | |
| TchangeHot | 0.479 | 0.155 | 22.977 | | | | |
| massScaled | 0.208 | 0.1 | 21.327 | | | | |
| **Type** | **Sum Sq** | **Mean Sq** | **NumDF** | **DenDF** | **F value** | **Pr(>F)** | |
| TchangeHot | 2.308 | 2.308 | 1 | 22.977 | 9.515 | 0.005 | |
| massScaled | 1.049 | 1.049 | 1 | 21.327 | 4.323 | 0.05 | |

##### 2.3 Table S3: Gene expression changes between ages in heat and cold

Table 3: The number of differentiall expressed genes that fall into each category of expression change across age, in response to acute heat (columns) and in response to acute cold (rows).

| | In heat | | | | | |
|---|---|---|---|---|---|---|
| | just_1week | just_8week | not_sig | opposite_direction | same_direction | same_slope |
| **In cold** | | | | | | |
| just_1week | 27 | 3 | 2 | 1 | 31 | 0 |
| just_8week | 11 | 2 | 4 | 2 | 8 | 0 |
| not_sig | 37 | 9 | 0 | 2 | 16 | 132 |
| opposite_direction | 8 | 1 | 1 | 7 | 20 | 0 |
| same_direction | 9 | 3 | 0 | 4 | 21 | 0 |
| same_slope | 0 | 0 | 11 | 0 | 0 | 44 |

"Not_sig" denotes genes that were not differentially expressed at the respective temperature.

#### 2.4 Table S4: Gene function over-representation analysis of model without age interaction

Table 4: Results of GO and KEGG over-representation analyses on the genes of the indicated response types (each gene is in only one row) with adjusted p-values < 0.1. Chicken annotations were used.

| **Condition** | **direction** | **Nr** | **GO** | **KEGG** |
|---|---|---|---|---|
| Hot | up | 25/32 | protein (re)folding, protein maturation, (cellular) response to unfolded protein, response to temperature stimulus | Spliceosome |
| hot | down | 112/148 | NA | Apoptosis, C-type lectin receptor signaling pathway (immune system) |
| cold | up | 2/4 | (neutral/cellular) lipid metabolic process; acylglycerol/glycerolipid metabolic process; (glycero-)lipid biosynthetic process | Glycerolipid metabolism, Cellular senescence |
| cold | down | 6/13 | cell migration, regulation of blood pressure, pyrimidine ribonucleotide metabolic process, multicellular organism development, phagocytosis | ECM-receptor interaction, Nucleotide metabolism, Integrin signaling, Biosynthesis of cofactors, Efferocytosis, Cytoskeleton in muscle cells, Focal adhesion |
| shared | hot up cold down | 7/10 | RNA processing, pseudouridine synthesis, fatty acid transport, negative regulation of (endo)peptidase activity, monocarboxylic acid transport | Mucin type O-glycan biosynthesis, lipid metabolism, Vascular smooth muscle contraction, Cytoskeleton in muscle cells |
| shared | hot down cold up | 14/15 | NA | Toll-like receptor & MAPK signalling pathways, Glycine, serine and threonine metabolism |
| shared | hot up cold up | 6/9 | organelle transport along microtubule, vesicle localization/trafficking, anterograde axonal transport, mononuclear & myeloid cell differentiation | Cytokine-cytokine receptor interaction, Lysosome biogenesis |
| shared | hot down cold down | 41/50 | multicellular organism development | Steroid hormone biosynthesis |

#### 2.5 Table S5: Gene function over-representation analysis of model with age interaction

Table 5: Results of GO and KEGG over-representation analyses with adjusted p-values $< 0.1$ on the genes that change temperature related gene expression between age groups. Chicken annotations were used. Numbers refer to the number of genes with available functional annotation out of the total number of genes in each category.

| Condition | Age.cat | Nr | GO | KEGG |
|---|---|---|---|---|
| Hot | Just 1 week | 65/92 | NA | NA |
| hot | Just 8 week | 10/18 | response to external biotic stimulus, cellular response to oxygen-containing compound, cell adhesion | Starch and sucrose metabolism, Cell adhesion molecule (CAM) interaction, Neuroactive ligand signaling, Regulation of actin cytoskeleton |
| hot | Same direction | 82/96 | NA | Calcium signaling pathway |
| hot | Opposite direction | 8/16 | NA | Terpenoid backbone biosynthesis |
| cold | Just 1 week | 50/64 | NA | NA |
| cold | Just 8 week | 21/27 | regulation of peptidase activity, regulation of cysteine-type endopeptidase activity involved in apoptotic process | NA |
| cold | Same direction | 26/37 | NA | NA |
| cold | Opposite direction | 29/37 | synaptic vesicle endocytosis, import into cell, potassium ion import across plasma membrane | Influenza A |
